# Continuous B- to A-Transition in Protein-DNA Binding - How Well Is It Described by Current AMBER Force Fields?

**DOI:** 10.1101/2022.01.13.476176

**Authors:** Petr Jurečka, Marie Zgarbová, Filip Černý, Jan Salomon

## Abstract

When DNA interacts with a protein, its structure often undergoes significant conformational adaptation. Perhaps the most common is the transition from canonical B-DNA towards the A-DNA form, which is not a two-state, but rather a continuous transition. The A- and B-forms differ mainly in sugar pucker P (north/south) and glycosidic torsion χ (high-*anti*/*anti*). The combination of A-like P and B-like χ (and *vice versa*) represents the nature of the intermediate states lying between the pure A- and B- forms. In this work, we study how the A/B equilibrium and in particular the A/B intermediate states, which are known to be over-represented at protein-DNA interfaces, are modeled by current AMBER force fields. Eight protein-DNA complexes and their naked (unbound) DNAs were simulated with OL15 and bsc1 force fields as well as an experimental combination OL15χ_OL3_. We found that while the geometries of the A-like intermediate states in the molecular dynamics (MD) simulations agree well with the native X-ray geometries found in the protein-DNA complexes, their populations (stabilities) are significantly underestimated. Different force fields predict different propensities for A-like states growing in the order OL15 < bsc1 < OL15_χOL3_, but the overall populations of the A-like form are too low in all of them. Interestingly, the force fields seem to predict the correct sequence-dependent A-form propensity, as they predict larger populations of the A-like form in naked (unbound) DNA in those steps that acquire A-like conformations in protein-DNA complexes. The instability of A-like geometries in current force fields may significantly alter the geometry of the simulated protein-DNA complex, destabilize the binding motif, and reduce the binding energy, suggesting that refinement is needed to improve description of protein-DNA interactions in AMBER force fields.

## Introduction

The conformational changes induced in DNA upon protein binding are linked to the affinity and selectivity of protein-DNA interactions and in turn to the function of protein-DNA complexes in DNA transcriptional regulation, processing, packaging and other protein-mediated DNA roles. Reliable theoretical (here, force field) modeling of protein-induced DNA adaptations requires accurate description of the conformational changes of the sugar-phosphate backbone, including sugar puckering and glycosidic angle, with particular attention to the relative energies of all energetically accessible conformers.

When a protein interacts with a DNA fragment, it can use a variety of mechanisms to find its target sequence, including base readout or shape readout.^1^ The shape of DNA and its changes accompanying complex formation are usually described in terms of helical parameters, groove widths, bend, etc. Underlying these helical changes are conformational changes at the atomistic level, such as sugar re-puckering, glycosidic bond rotation, BI/BII flipping or many other sugar-phosphate backbone reconformations. In the following we will focus on this backbone response to DNA complexation by a protein. While detailed backbone reconformations are less frequently discussed than the global helical parameters of the duplex, from the point of view of the force field modeling, the backbone angle shifts or flipping is the primary cause of the change in the global helical structure.

Deformations from canonical B-DNA towards the A-DNA form are among the most common conformational adaptations of DNA interacting with a protein. The protein-induced transition to the A-form may be incomplete, featuring structures intermediate between the A- and B- forms.^2, 3^ This type of structural adaptation is very common in various unrelated protein-DNA complexes, where, for example, helices of zinc finger, beta sheet or helix-turn-helix structures interact with the major groove, leading to its widening.^4^ The extent of the B->A conformational conversion and related structural motifs in a number of protein-DNA complexes have been studied by the Olson’s group^5^ and it has been suggested that the propensity to adopt the A-form in a complex follows the scale of A-forming tendencies observed for naked DNA duplexes in solution, as described by Ivanov et al.^6^

As mentioned above, DNA conformational adaptations are usually described in terms of groove widths, DNA helical parameters, such as twist, slide or inclination, or the *Z*p parameter.^7^ These descriptors do not provide immediate information about the conformation of the sugar-phosphate backbone and it is not straightforward to transform changes in helical parameters into the underlying backbone conformations. However, backbone conformations are of utmost importance for molecular modeling, because they are the native language of atomistic empirical force fields. Backbone conformations upon protein to DNA binding have been studied in detail by Schneider et al.^8^ The induction of the A-DNA form during complex formation has been attributed to A-like and intermediate A/B conformers, which are rare in naked DNA duplexes. Importantly, the intermediate A/B conformers were characterized by the average values of the backbone dihedral angles and glycosidic angle determining the detailed atomistic structures of the intermediate A/B states. The conformations of A-DNA, B-DNA and selected midway conformations are shown in Figure 1.

**Figure 1.**
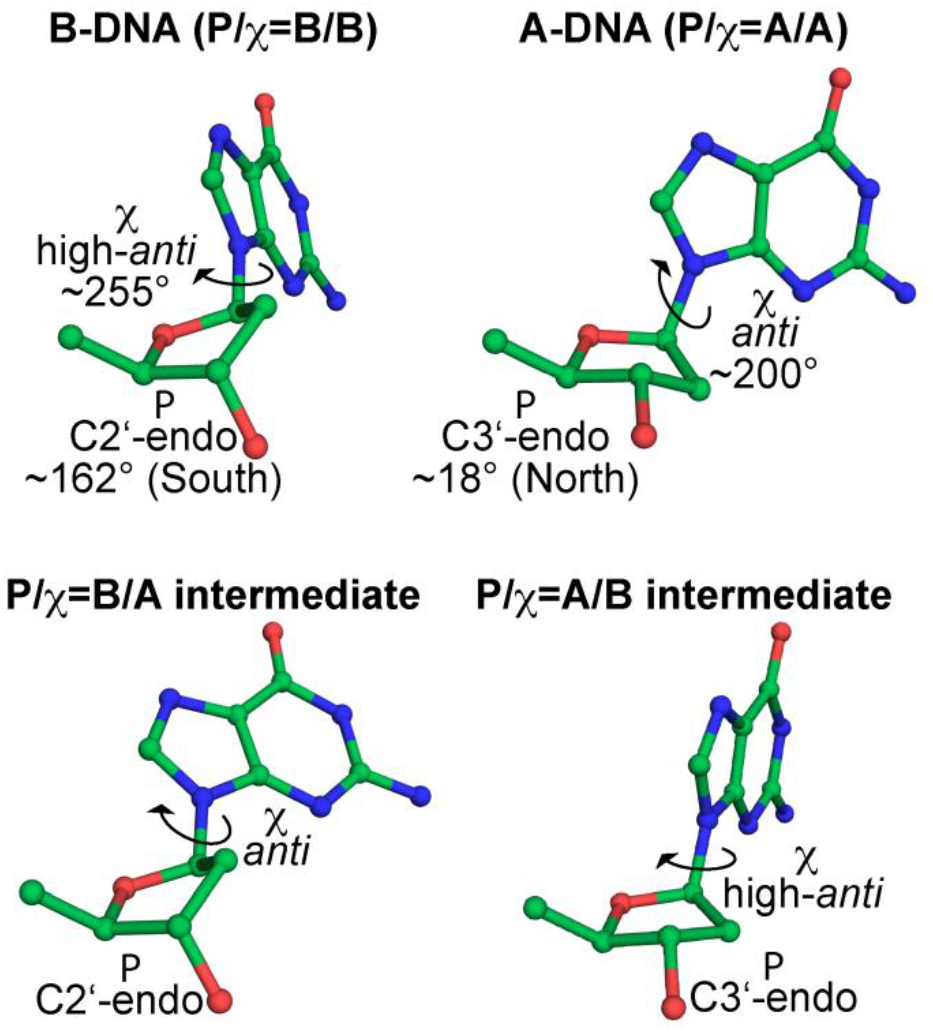
Sugar pucker and glycosidic torsion χ in A-DNA, B-DNA and two common intermediate conformers frequently found in protein-DNA complexes, present in dinucleotide conformations AA02 and BB16 (see ref. ^9^).

The aim of this work is to investigate how well the conformational landscape of the deoxyribose pucker and glycosidic angle is described by current well established DNA force fields from the AMBER family of force fields. We note that deoxyribose puckering is also debated and important topic in the CHARMM force field,^10,11^ the recent AMOEBA force field for nucleic acids^12^ and a recent version of the AMBER force field that modifies, among other, the puckering parameters.^13^ Currently, there are two widely used modifications, our OL15^14^ and bsc1^15^ from the Orozco group in Barcelona. (Note that a new version of the OL force field, OL21^16^ has recently been published, but it does not modify the sugar pucker or χ and testing it would therefore not provide useful additional information in this work.) Both mentioned force fields provide a reasonably accurate description of the basic B-DNA characteristics,^17–19^ however, both have similar shortcomings in describing the A/B equilibrium in naked DNA in water and in ethanol (or trifluoroethanol) solutions.^20^ The fact that neither our OL15 nor bsc1 correctly predicts A-DNA stability in solution led us to investigate A/B equilibrium in protein-DNA complexes, and as we show below, the problems observed in naked DNA actually manifest themselves in the interactions of DNA with protein.

OL15 and bsc1 were introduced in 2015 and both are based on the same parent force field ff99^21,22^ and introduced several modifications to backbone dihedral angles (Table 1). As can be seen from the table, both OL15 and bsc1 modified the glycosidic torsion angle χ (the χ modification used in OL15 comes from earlier work^23^) and also the ε and ζ potentials (ε/ζ modification used in OL15 comes from our 2013 work^24^). However, OL15 also modifies also the β potential,^14^ which was left unchanged in bsc1. This β modification improved the description of non-canonical Z-DNA, but also the BI/BII equilibrium in the B-DNA diplex.^14^ Furthermore, while the sugar pucker parameters were not modified in OL15, they were changed in the bsc1 force field.^15^ In addition, further testing is performed on modified OL15, OL15χ_OL3_, in which its χ_OL4_ glycosidic parameterization^23^ was replaced by χ_OL3_ parameterization developed earlier for RNA.^25^ This is motivated by the desire to estimate the effect of χ parameterizaion on the results (note that χ_OL3_ was derived for A-RNA and may provide a better description of the A-like region).

**Table 1.** An overview of the DNA force fields used.

| Combination | parent FF | $\alpha/\gamma$ | $\beta$ | $\epsilon/\zeta$ | $\chi$ | Pucker |
| --- | --- | --- | --- | --- | --- | --- |
| OL15 | parm99<sup>22</sup> | bsc0<sup>26</sup> | $\beta_{OL1}$<sup>14</sup> | $\epsilon/\zeta_{OL1}$<sup>24</sup> | $\chi_{OL4}$<sup>23</sup> | parm99 |
| OL15$\chi_{OL3}$ | parm99 | bsc0 | $\beta_{OL1}$ | $\epsilon/\zeta_{OL1}$ | $\chi_{OL3}$<sup>25</sup> | parm99 |
| bsc1 | parm99 | bsc0 | parm99 | bsc1 $\epsilon/\zeta$<sup>15</sup> | bsc1 χ<sup>15</sup> | bsc1 P<sup>15</sup> |

In the following, we discuss the stability of A-DNA conformations and A/B intermediate conformations in protein-DNA complexes described by OL15, OL15χ_OL3_ and bsc1 force fields.

## Methods

### Selection of Protein-DNA Complexes

Protein-DNA structures were selected to contain well-defined representatives of B-DNA, A-DNA and also structures intermediate between A- and B-DNA according to the NtC dinucleotide classification of Černý et. al.^9^ We focused on two representative A/B intermediate nucleotide structures, present in AA02 and BB16 NtC dinucleotides. AA02 dinucleotide contains two consecutive sugars that have A-type (North, C3’-endo) sugar puckering but B-type glycosidic torsion χ (high-*anti*, ~245°); one of these nucleotides is shown in Figure 1. BB16, on the other hand, has a nucleotide with a B-type (South, C2’-endo) sugar puckering and an A-type glycosidic torsion χ (*anti*, ~205°); nucleotide with this conformation is also shown in Figure 1. In this regard, AA02 and BB16 represent two clear examples of structures that are intermediate between the A- and B- forms. They are also quite frequently found in X-ray databases^9^ and AA02 has been shown to be over-represented at the protein-DNA interface compared to free DNA (see cluster 32 in ref.^8^). Note that there are other types of A/B intermediate structures with P or χ on the way between A- and B-forms. Some of these are also present in the selected protein-DNA complexes and were included in our analysis.

In our selection, we focused on structures that contained well-defined NtC dinucleotide representatives AA00 (pure A-DNA), AA02 and BB16 in that the structures had good resolution and the NtCs had high “confal” scores, which quantify the conformity between the analyzed dinucleotide and the reference NtC geometry.^27^ Different types of protein-DNA complexes were selected in terms of protein binding motifs and their localization to major/minor grooves, with the exception of two structures of chromatin-DNA complexes (1C8C and 3KXT), which are quite similar and served as a test of whether our MD simulations can provide consistent results for similar complexes. The eight selected complexes are listed in Table 2 and shown in Figure 2. Because we are interested in the behavior of single nucleotide units rather than dinucleotides, which are used in the NtC classification, we focus in the following on the analysis of the sugar pucker P and glycosidic torsion χ of individual nucleotides rather than on the dinucleotide NtC classes.

**Table 2.** Selected protein-DNA complexes. Terminal bases not included in the number of states.

| PDB ID | DNA complex with | Resolution | Number of states with $P/\chi =$ | | | |
| --- | --- | --- | --- | --- | --- | --- |
|  |  |  | A/A | A/B | B/A | B/B |
| 1DP7<sup>28</sup> | hRFX1 factor | 1.50 | 6 | 0 | 0 | 22 |
| 1P71<sup>29</sup> | HU architectural factor | 1.90 | 2 | 2 | 3 | 29 |
| 1QNE<sup>30</sup> | TATA box binding prot. | 1.90 | 1 | 9 | 0 | 14 |
| 2VJV<sup>31</sup> | TnpA transposase | 1.90 | 19 | 2 | 5 | 30 |
| 3AAF<sup>32</sup> | RecQ helicase | 1.90 | 4 | 1 | 1 | 18 |
| 3PVI<sup>33</sup> | PvuII endonuclease | 1.59 | 17 | 0 | 0 | 5 |
| 1C8C<sup>34</sup> | Sso7d chromatin prot. | 1.45 | 0 | 5 | 0 | 7 |
| 3KXT<sup>35</sup> | Cren7 chromatin prot. | 1.60 | 0 | 3 | 0 | 9 |

**Figure 2.**
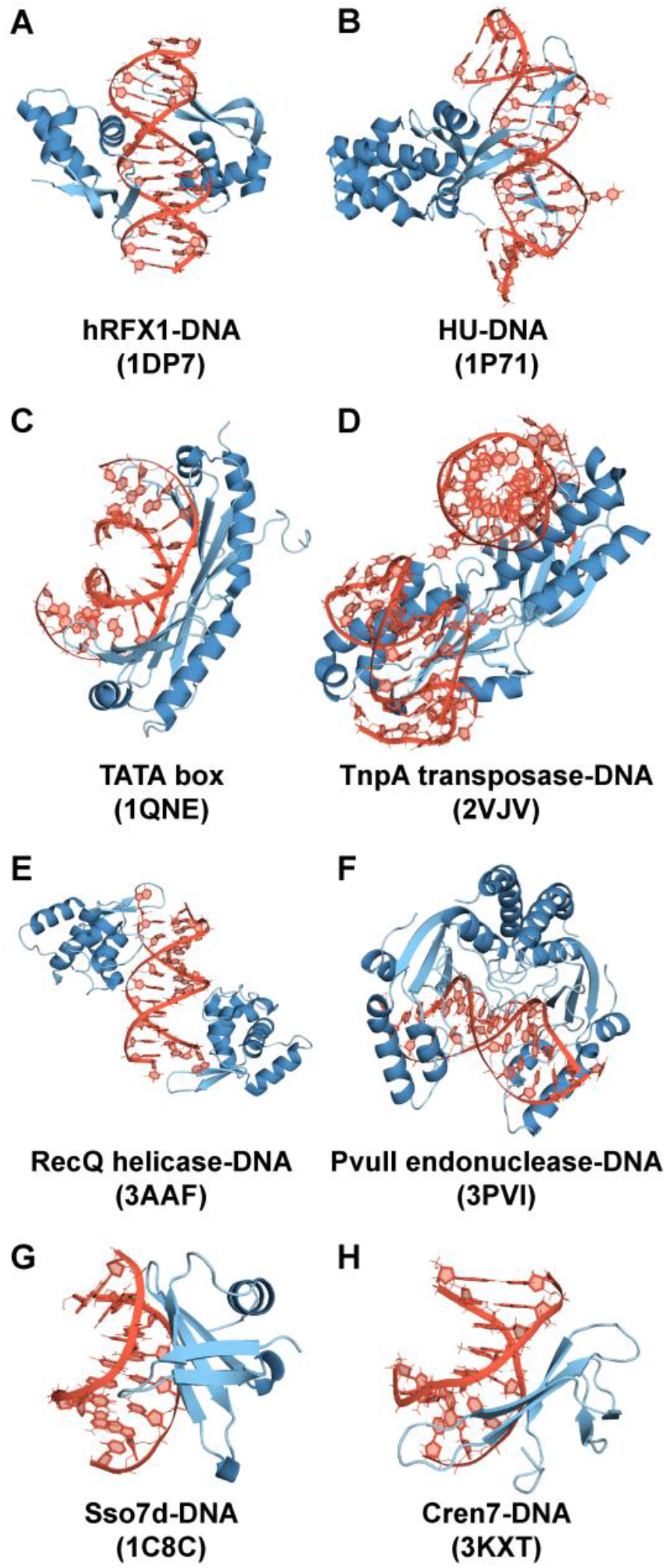
Protein-DNA complexes studied in this work (PDB ID in the parentheses).

### MD Simulations

Table 1 details the force field combinations used in this work. The initial structures listed in Table 2 were first neutralized with K^+^ cations and the ion-concentration was then adjusted to 0.15M KCl using ionic parameters by Joung and Cheatham.^36, 37^ The complexes were solvated with an octahedral box with a 10 Å buffer and water molecules identified in the X-ray structures were kept.

Relaxation of the initial structures was performed as described previously.^38^ Molecular dynamics simulations were performed using the CUDA PMEMD code from AMBER 18^39^ under NPT conditions (1bar, 298 K) with Monte Carlo barostat pressure control (taup=2), Langevin thermostat, and a time step of 4 fs, using the hydrogen mass repartitioning procedure.^40, 41^ 10 Å direct space nonbonded cutoff and SHAKE on bonds on hydrogen atoms with default tolerance (0.00001 Å) were used and the nonbonded pair list was updated every 25 steps. Coordinates of the protein, nucleic acid and ions were stored every 10 ps.

When Mg^2+^ ions were present in the crystal structure, they were kept and described by parameters of Allnér et al.^42^ BrU residues were replaced by dT residues in 1DP7. When protein chain was not continuous (missing residues) the ends were terminated with ACE and NME residues. Protonation states of histidine residues were assigned using the PropKa server,^43^ and corrected when necessary based on visual inspection of the hydrogen bonding network. Simulations of free DNA molecules (without protein) were initiated from the geometry in protein-DNA complex.

The analysis of nucleic acid structure parameters was performed using the cpptraj module and nastruct tool of AMBER software package.

### Classification of Backbone Conformations

Sugar pucker P was considered North (N) if P was less than 90° or greater than 270°, otherwise it was considered South (S). The glycosidic angle χ was considered *anti* in the interval between 120° and 227.5° and high-*anti* when it was greater than 227.5° (*syn* otherwise; *syn* populations were negligible and are not reported). The value of 227.5 was chosen as a midpoint between the χ values for pure B-form (BB00) and pure A-from (AA00) according to Černý et al.^9^ This distinction is arbitrary and it should be kept in mind that the A- and B-like χ distributions are relatively wide and may overlap, therefore a clear and unambiguous distinction is not possible.

## Results and Discussion

### Overall Stability of Simulated Complexes

Most simulations were relatively stable in all tested force fields. The RMSDs of the whole complex, nucleic acid and protein are shown separately in the Supporting Information Figure S1. Larger deviations can be seen, for instance, in the OL15χ_OL3_ simulation of the chromatin-DNA complex 1C8C, which is due to fraying of the terminal base pairs. In general, some amount of fraying can be expected in most of our simulations because the DNA helix ends are often stacked on its neighboring molecules in the crystal structure and are thus stabilized by packing, whereas in the simulations they are free. The larger deviation in the bsc1 simulation of 1C8C is due to one of the central AT pairs losing pairing. All three force fields give relatively large RMSD for the nucleic acid of the HU-DNA architectural factor, but this is likely due to the duplex being bent significantly due to crystal contacts in the X-ray structure and following relaxation in the MD simulations. Notably, the larger RMSD for endonuclease 3PVI is mainly due to the extensive transition from the A-like form towards the B-DNA form, which is more pronounced in the OL15 simulation. The somewhat larger RMSD for helicase 3AAF has the same cause, but its magnitude is smaller due to the lower native content of the A-form in the crystal. Overall, the three force fields perform similarly and the small differences are likely random and statistically insignificant.

### Distributions of P and χ in Intermediate States

The distributions of P and χ in a canonical B-DNA nucleotide, an A-DNA nucleotide and two A/B and B/A intermediate nucleotides are shown in Figure 3. The data are taken from the OL15 MD simulations of the chromatin-DNA complex 1C8C and transposase 2VJV, where we selected four representative nucleotides that retain their native state throughout the simulation: dC106_B and dC114_C in 1C8C (B/B and A/B states, respectively) and dT29_D and dC20_C in 2VJV (B/A and A/A states, respectively). Vertical lines indicate reference values of χ taken from Černý et al.^9^ (NtCs BB00, AA00, AA02 and BB16) and sugar pucker values are averages over all residues in a given state in our eight complexes (note that P values are not provided in the database by Černý et al.).

**Figure 3.**
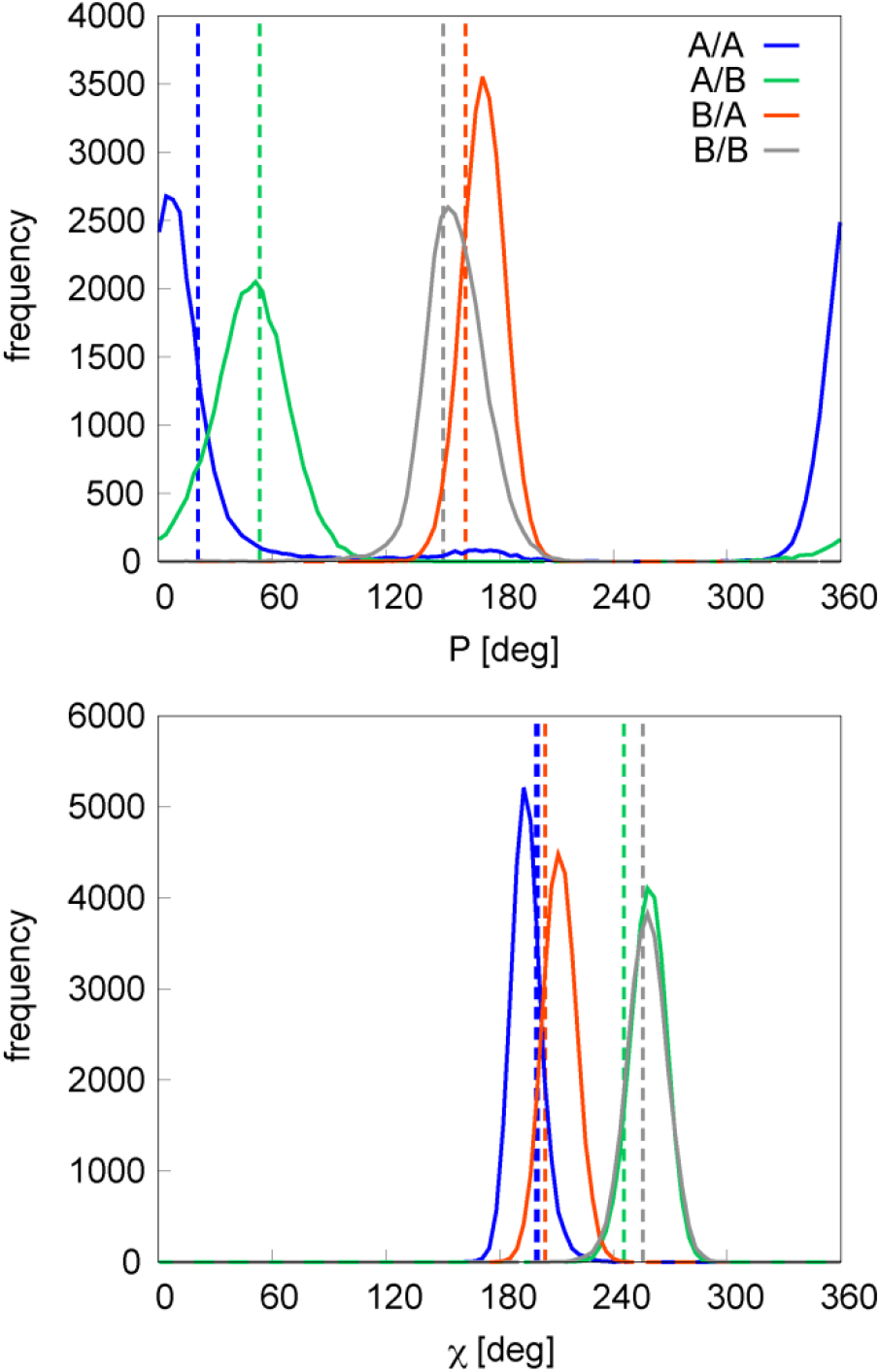
Distributions of P and χ in nucleotides representative for B/B (dC106_B, 1C8C), A/B (dC114_C, 1C8C), B/A (dT29_D in 2VJV) and A/A (dC20_C, 2VJV). Dashed vertical lines represent reference values from X-ray (see text).

As can be seen from the average values for our protein-DNA X-ray geometries (vertical lines in Figure 3), the sugar pucker in the pure P/χ = A/A form (21°) differs significantly from the pucker in the intermediate A/B form (53°). The pucker of the intermediate A/B form is thus on the way between the pure A/A and B/B forms. Similarly, χ of the intermediate A/B and B/A forms lies between the pure A/A and B/B forms, although always much closer to its parent. Interestingly, opposite is true for the sugar pucker in the B/A intermediate form, which is shifted away from the A/A value and is thus even larger than that of the pure B/B form. When we compare these X-ray values with the distributions from the OL15 MD simulations, we can see that the trends described above hold and the MD distributions are close to the respective X-ray values. The selected nucleotides are quite representative in this respect, and similar trends are seen for the other nucleotides that retained their native P/χ configurations in the simulations. Thus, the geometries of the A/B intermediate states are modeled reasonably well by the OL15 force field.

### Time Development of A-like Backbones in MD Simulations

Different nucleotides exhibit different behaviors of the A-like backbone conformations in the MD simulations. In the following we comment on several possible behaviors for each of the studied states.

### P/χ = A/A States

In complexes where P/χ = A/A states are highly populated during the simulation, like for dA8 in 1DP7 factor X complex (OL15), the B-like (S) pucker is frequently visited during the simulation, but is unstable and the conformation quickly reverts to the A-like (N) state. Short visits to the S pucker state are accompanied by small shifts of the χ angle from the A-like *anti* region towards the high-*anti* region, but higher values are only reached during the longest pucker reconformations (Figure 4). In some cases, the A/A form is stable for a few tens of ns but then converts to the B/B form, as in the case of dC7 residue in Figure 4. Another example is residue dT9 in the same structure, which transitions from the A/A form to the B/B form very quickly, within the first ns of the simulation and show no tendency to return to the initial A/A state. This is the case for most residues in the 3PVI endonuclease and 2VJV transposase complexes. These three types of behavior are also found in other simulated structures.

**Figure 4.**
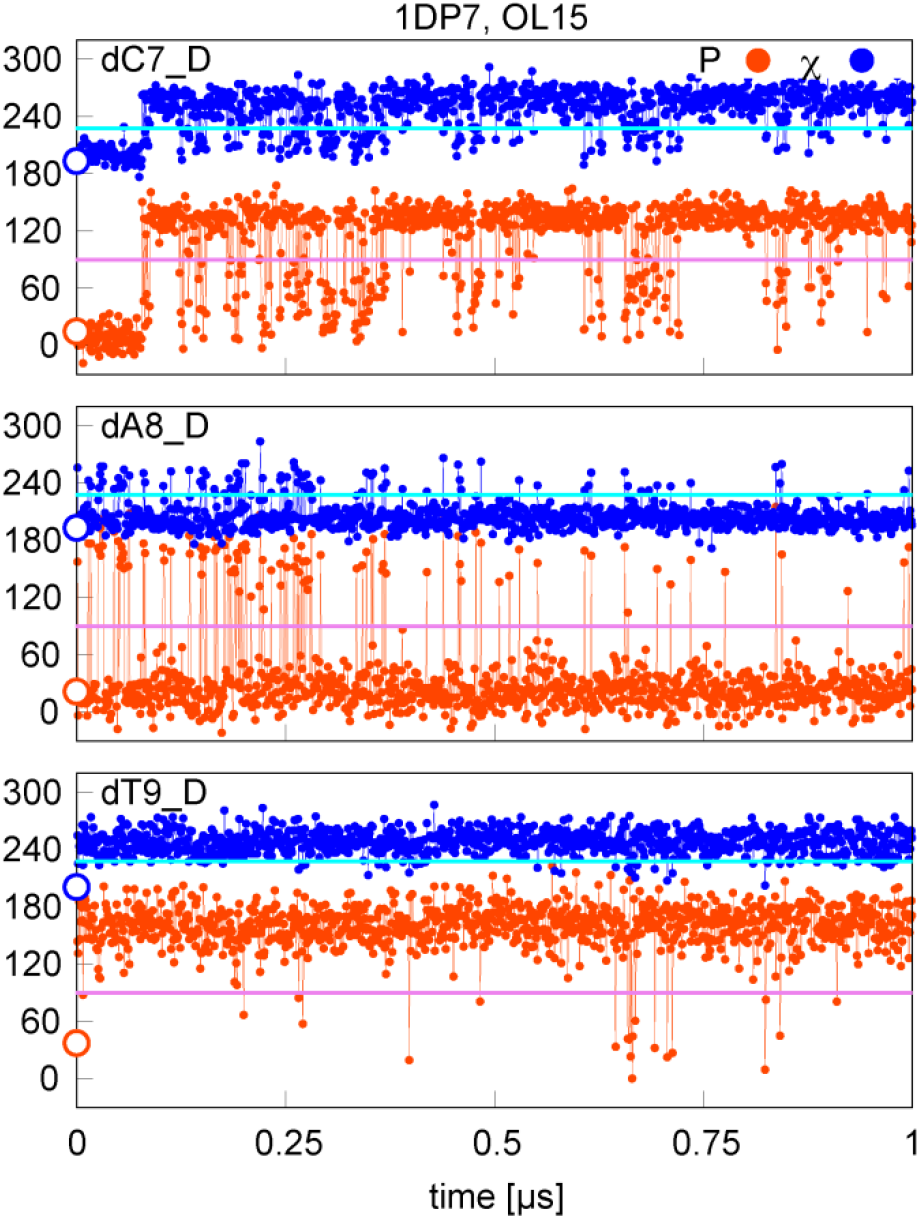
Sugar pucker and χ in OL15 simulations of selected residues with starting (native) P/χ = A/A state in the 1DP7 complex. Values separating A- and B-type P and χ are shown in pink and cyan, respectively; points on the y-axis indicate X-ray values.

### P/χ = A/B States

In case of P/χ = A/B states, if they are stable, as is the for residue dT113 of the chromatin complex 3KXT (OL15), the sugar pucker sometimes visits the B-like region, but this visit is transient and the pucker quickly returns to the A-like state (Figure 5). Another possible behavior is that the sugar pucker again remains in the A-like state, but χ is on the borderline between the A- and B-forms, leading to a partial assignment to the A/A state, as in residue dT105 of 3KXT in Figure 5. This example represents only a minor deviation from the initial A/B state, which is conditional on the (arbitrary) choice of the χ division line, but can still indicative of the accuracy of the force field used; OL15 is closest to the experimental A/B geometry, with bsc1 and OL15χ_OL3_ being somewhat more distant. Another case is the dG103 residue in 3KXT, where the sugar often visits the B-like pucker, leading to well defined B/B states.

**Figure 5.**
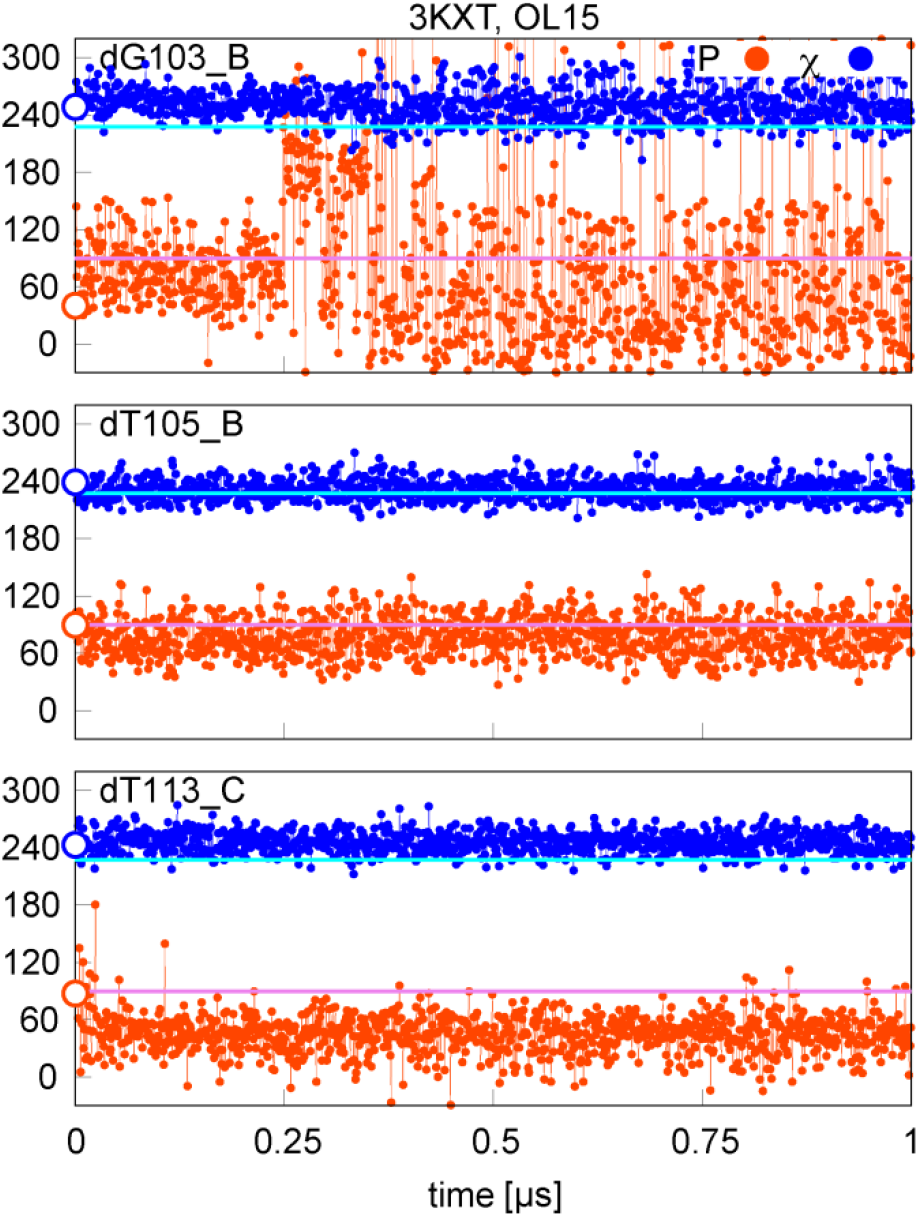
Sugar pucker and χ in OL15 simulations of selected residues with starting (native) P/χ = A/B state in the 3KXT complex. Values separating A- and B-type P and χ are shown in pink and cyan, respectively; points on the y-axis indicate X-ray values.

### P/χ = B/A States

The P/χ = B/A states are often stable (but not fully) in all tested force fields. When they deviate from the B/A conformation, it is usually because the χ angle visits the B-like (high-*anti*) region as in residue dT35 in 2VJV (Figure 6). This is very frequent, because of the anti/high-anti overlap, and in fact it is not possible to clearly distinguish the B/A and B/B forms because of a relatively small shift in the χ angle. Here, bsc1 is somewhat better than OL15 and OL15χ_OL3_ provides results closest to the X-ray data. A less frequently observed behavior is the visiting of the A-like pucker region as in residue dT30 in chain D of the 2JVJ structure, which is, however, not very populated.

**Figure 6.**
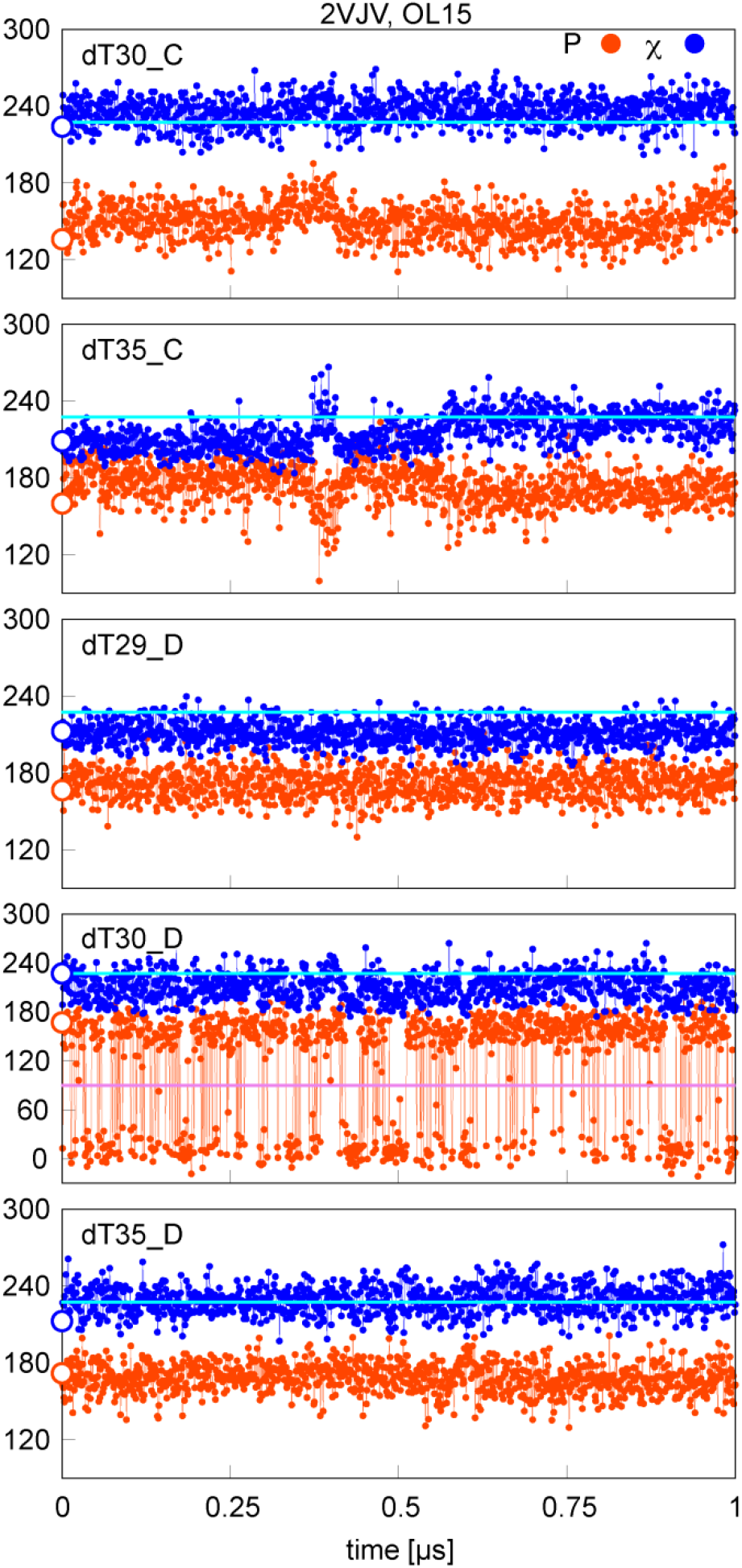
Sugar pucker and χ in OL15 simulations of selected residues with starting (native) P/χ = B/A state in the 2VJV complex. Values separating A- and B-type P and χ are shown in pink and cyan, respectively; points on the y-axis indicate X-ray values.

### Average Stability of A-like Backbones

As described above, the time evolution of P and χ in MD simulationsan can vary significantly for individual residues in different complexes. Nevertheless, it may be informative to consider MD populations of individual P/χ conformers averaged over multiple nucleotides starting in the same initial (native) state. Table 3 shows the percentages of native X-ray P/χ states retained in simulations of individual protein-DNA complexes for all tested force fields. For instance, in 1DP7 complex, only 30% of the initial A/A conformations and 91% of the initial B/B conformations were present in the OL15 simulation (there were no A/B or B/A conformations in the X-ray structure of 1DP7).

**Table 3.** Percentages of P/χ = AA, AB, BA and AB states in X-ray and in MD simulations. Terminal residues were excluded.

| PDB ID | % of native X-ray $P/\chi$ states preserved in MD simulations | | | | | | | | | | | |
| --- | --- | --- | --- | --- | --- | --- | --- | --- | --- | --- | --- | --- |
| | OL15 | | | | bsc1 | | | | OL15$\chi_{OL3}$ | | | |
|  | A/A | A/B | B/A | B/B | A/A | A/B | B/A | B/B | A/A | A/B | B/A | B/B |
| 1DP7 | 30 | - | - | 91 | 61 | - | - | 90 | 60 | - | - | 83 |
| 1P71 | 12 | 78 | 46 | 79 | 13 | 64 | 65 | 78 | 16 | 72 | 61 | 70 |
| 1QNE | 40 | 95 | - | 74 | 63 | 91 | - | 70 | 74 | 91 | - | 50 |
| 2VJV | 37 | 85 | 59 | 84 | 41 | 70 | 66 | 81 | 47 | 75 | 70 | 72 |
| 3AAF | 1 | 0 | 14 | 89 | 2 | 0 | 17 | 90 | 4 | 1 | 17 | 80 |
| 3PVI | 5 | - | - | 91 | 3 | - | - | 87 | 16 | - | - | 76 |
| 1C8C | - | 52 | - | 97 | - | 35 | - | 89 | - | 47 | - | 77 |
| 3KXT | - | 64 | - | 88 | - | 54 | - | 83 | - | 50 | - | 72 |

It can be seen From Table 3 that the stability of various A-like states varies between different types of protein-DNA complexes. For instance, in the helicase 3AAF, endonuclease 3PVI and architectural factor 1P71 complexes the A-like structures (P/χ = A/A) were mostly lost very rapidly, often within the first ns of the simulation. In contrast, a high degree of A-form preservation is observed in some complexes where the DNA is significantly distorted by the protein. For example, in the TATA box 1QNE the β-sheet of the protein interacts with the minor groove, which is subsequently significantly widened, bent and the helical twist is strongly reduced in this region. All residues found in the 1QNE crystal in the P/χ = A/B intermediate form retain this conformation in all our simulations. The situation is similar, to a lesser extent, in the chromatin proteins 1C8C and 3KXT, where the minor groove is again widely opened by the β-sheet. Also for the 1P71 cofactor, the interaction with a loop/β-sheet structure widens the minor groove and the P/χ = A/B structures are well preserved in our simulations. Nevertheless, the population of the A-like native structures appears to be too low in most of the simulated structures.

The populations averaged over all simulated structures are given in Table 4. In this table we also list the populations of the non-native states in MD simulations to which the original (native) state is converted in the MD simulation. The populations of native states (conservation percentages) corresponding to the values in Table 4 are highlighted in bold.

**Table 4.**
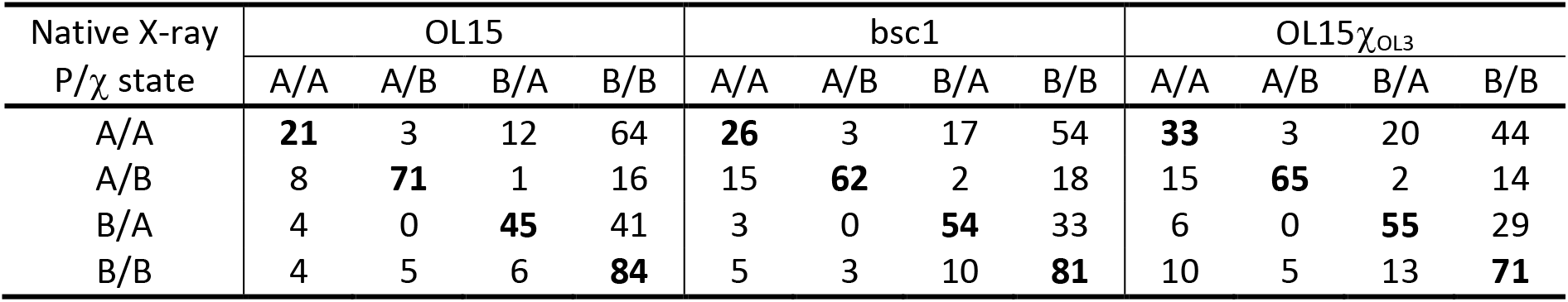
Percentage of P/χ = AA, AB, BA and AB states in MD simulations averaged over all structures, grouped by the native (starting) X-ray state. Populations of native states are highlighted in bold. Terminal residues were excluded.

| Native X-ray<br>$P/\chi$ state | OL15 | | | | bsc1 | | | | OL15$\chi_{OL3}$ | | | |
| --- | --- | --- | --- | --- | --- | --- | --- | --- | --- | --- | --- | --- |
|  | A/A | A/B | B/A | B/B | A/A | A/B | B/A | B/B | A/A | A/B | B/A | B/B |
| A/A | **21** | 3 | 12 | 64 | **26** | 3 | 17 | 54 | **33** | 3 | 20 | 44 |
| A/B | 8 | **71** | 1 | 16 | 15 | **62** | 2 | 18 | 15 | **65** | 2 | 14 |
| B/A | 4 | 0 | **45** | 41 | 3 | 0 | **54** | 33 | 6 | 0 | **55** | 29 |
| B/B | 4 | 5 | 6 | **84** | 5 | 3 | 10 | **81** | 10 | 5 | 13 | **71** |

In general, the populations of A-like states seem to be too small in many cases in Table 3 and Table 4. We did not expect 100% conservation of all native P/χ configurations in our MD simulations, because even a small energy advantage over other states can lead to “freezing” of a more stable (more populated) state in the crystal structure. However, it seems unlikely that a state whose population is significantly less than 50% in the MD simulation (such as the A/A states in 3AAF) would emerge as dominant in the crystal structure. A more plausible explanation is that the stability of the A-like states is underestimated by the tested force fields. This is supported by the marked differences between the force field variants, which can be explained by varying quality of the description of the A-like structures (the tested force fields differ, among other, in the sugar pucker and χ potentials). The underestimation of the stability of the A-like states is also consistent with the results of our previous work, showing that OL15 and bsc1 force fields underestimate the stability of A-DNA in aqueous and ethanol solution.^20^

However, there are notable differences in the stability of the A/A, A/B and B/A conformers. The least stable is the P/χ = A/A conformer, the most common A-DNA form, with an average population of less than 34% in all force fields tested (Table 4). Most striking is the very low A/A content in the MD simulations of the endonuclease 3PVI, helicase 3AAF and transposase 2VJV complexes (Table 3). In most cases, the initial (native) A/A conformation rapidly transformed to the B-DNA conformation, often within the first 1 ns of the simulation. The speed of the conversion also suggests that the used force fields predict an overly destabilized A-DNA form.

The second least stable non-canonical conformer is P/χ = B/A, whose population varies between 45 and 55% among the force fields tested. The stability of this state with respect to the native P/χ = B/B conformation is mainly influenced by the χ potential. From the lower population of the P/χ = B/A conformer it would appear that the χ potential over-favors the B-like form over the A-like form. At the same time, however, the B/A population appears to be somewhat too high in the OL15χ_OL3_ simulations of nucleotides with a native B/B state. Note also that the P/χ = B/A states cannot be very clearly distinguished from the B/B states (see also the separation line Figure 6). In addition, B/A is also the least populated conformer in our complexes, with only 9 occurences, all found in 3 complexes. Thus, although the χ potential clearly affects the relative stabilities of both the B/A–B/B and A/A–A/B structures, we can neither infer nor rule out that it requires a significant modification in future.

The P/χ = A/B conformer is the most stable of the non-canonical conformers, ranging between 62 and 71% in MD simulations with different force fields. Although a small additional stabilization of this geometry could be beneficial, the magnitude of this stabilization is clearly less than that required by the A/A conformer. Note that the sugar pucker value of this conformer is different from that of the pure A/A conformer (53° vs. 21°, respectively), and therefore its stability is not necessarily related to the (in)stability of the A/A state discussed above.

Taken together, it seems that the A-like forms could be stabilized either by further lowering of the *anti* (A-like) region of the χ potential or the C3’-region of the sugar pucker. The stability of the χ = *anti* region increases in order OL15 < bsc1 < OL15χ_OL3_ in our simulations and the results obtained with the most stabilizing OL15χ_OL3_ combination suggest that the P/χ = B/A might be on the verge of being overpopulated in naked DNA simulations, while the stability of the P/χ = A/A state is still too low. This suggests that it is the C3’-region of the sugar pucker potential that could provide the desired stabilization of the A-like forms.

### P/χ Conformers in Naked DNA

Because the non-canonical P/χ conformers can also be found in the simulations of free DNA duplexes in solution (albeit in small amounts), we performed simulations of naked DNAs taken from our protein-DNA complexes. The populations of the P/χ states in complex and in free DNA simulation with the OL15 force field are compared in Table 5. The results for bsc1 and OL15χ_OL3_ were similar and are shown in Supporting Information in Tables S2 and S3.

**Table 5.**
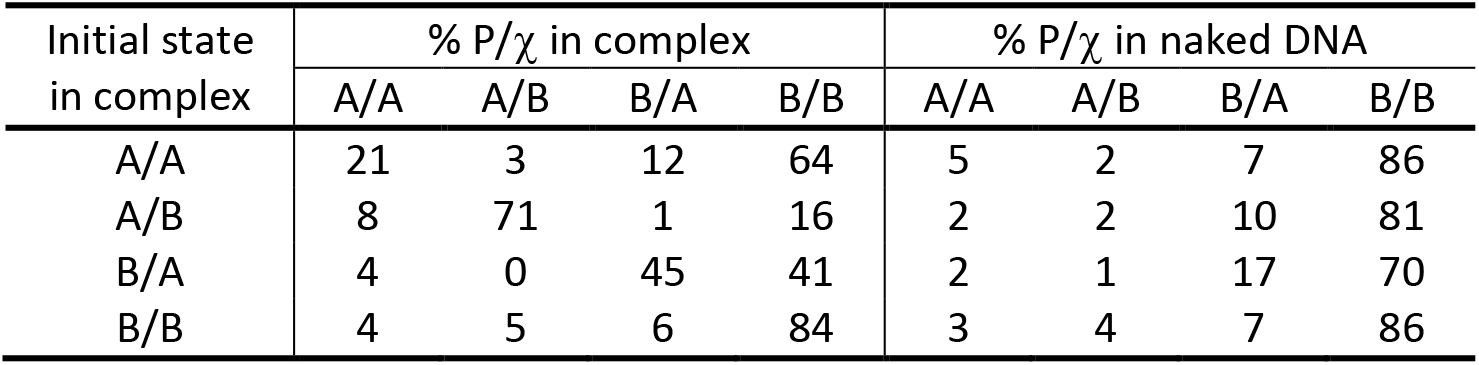
The propensity for different P/χ states in complexes and in their naked DNA averaged over all OL15 simulations.

| Initial state<br>in complex | % $P/\chi$ in complex | | | | % $P/\chi$ in naked DNA | | | |
| --- | --- | --- | --- | --- | --- | --- | --- | --- |
|  | A/A | A/B | B/A | B/B | A/A | A/B | B/A | B/B |
| A/A | 21 | 3 | 12 | 64 | 5 | 2 | 7 | 86 |
| A/B | 8 | 71 | 1 | 16 | 2 | 2 | 10 | 81 |
| B/A | 4 | 0 | 45 | 41 | 2 | 1 | 17 | 70 |
| B/B | 4 | 5 | 6 | 84 | 3 | 4 | 7 | 86 |

The propensity to the A-form is known to be sequence-dependent in DNA and it has been argued^5^ that the deformation towards the A-form in protein-DNA complexes follow the scale of the A-DNA propensities known from free DNA duplexes.^6^ This argument appears to be supported by our data, assuming that the force fields predict the correct relative sequence-dependent A-form propensities in free DNA simulations. Table 5 shows that the nucleotides that adopt A/A conformations in the protein-DNA complexes studied also show increased percentage of the A/A form in the naked DNA simulations (5%, see Table 5). Consistent with this, nucleotides that assume intermediate B/A conformations in our protein-DNA complexes are more prone to the B/A conformation in free DNA duplexes. Interestingly, this does not seem to be the case for the intermediate A/B structures. For instance, the steps that assume the A/B conformation in the protein/DNA complexes exhibit very low A/B population in free DNA simulations.

## Conclusions

The stability of A-like nucleotide conformations was investigated in force field simulations of protein-DNA complexes. Particular attention was paid to conformations intermediate between A-DNA and B-DNA forms, which are known to be overrepresented on protein-DNA interfaces compared to naked DNA simulations. Two established AMBER force fields, OL15 and bsc1, and one combined force field, OL15χ_OL3_, in which χ parameters are taken from the OL3 RNA force field, were tested. We found that when the A-like backbone substates are stable in the simulations, their geometry is described relatively well by the force field and reflects the average X-ray geometries known form the crystal structures. This applies not only to the pure A-DNA form (P/χ = A/A), but also to the conformations intermediate between A- and B-DNA, i.e., P/χ = A/A and P/χ = A/A, which are typical for protein-DNA complexes. This means that the force fields are capable of modeling the geometry of the continuous transition from B-DNA to A-DNA form during protein-DNA binding.

However, while the geometries of the A-DNA and intermediate A-like structures seem to be described well, their stabilities are probably underestimated by current AMBER force fields. This is most pronounced for pure A-DNA nucleotides (P/χ = A/A), whose populations are less than 50% for most of the simulated nucleotides. The stability of A-like forms increases in the order OL15 < bsc1 < OL15χ_OL3_, but average percentage conservation of for instance the native A/A states in the MD simulations is only 21, 26 and 33% on our 1 μs time scale. The population of the intermediate P/χ = A/B conformer with A-like sugar pucker and B-like χ is also underestimated, and also the P/χ = B/A conformer is not fully stable in all simulated cases. We attribute this mainly to inaccurate description of the sugar puckering and possibly (to a lesser extent) the glycosidic torsion in current force fields.

The instability of the A-like geometries varies among different protein-DNA complexes. In some complexes, the A-like geometry is lost within the first nanoseconds of simulation and the structure rapidly transitions to the canonical B-DNA duplex, changing the geometry of the complex, which is reflected by an increased in RMSD. In other cases, especially when the protein-induced DNA deformation is relatively large, such as when the protein widely opens the minor groove, many A-like residues are able to retain their native geometry in the simulation. However, even in these cases, overly unstable A-like geometries enforced by the protein can be expected to destabilize the binding motif and significantly reduce the binding energy. The instability of the A-like forms in the current AMBER force fields represents a serious problem for accurate modeling of protein-DNA interactions and modification of the description of the sugar puckering may be needed.

By comparing the content of A-like conformers in protein-DNA complexes and in MD simulations of naked (unbound) DNA of these complexes, we found that the force fields appear to predict the correct sequence-dependent propensity to A-form, as they predict larger populations of the A-like form in MD simulations of naked (unbound) DNA in those steps that acquire A-like conformations in protein-DNA complexes.

## Acknowledgements

This work was supported by Grant 20-28231S (P.J., M.Z.) from the Grant Agency of the Czech Republic.

## Supporting Information

The Supporting Information is available free of charge on the ACS Publications website at DOI:RMSD plots for all simulated complexes and populations of P/χ conformers in simulations of naked DNA duplexes for bsc1 and OL15χ_OL3_ force fields.

