## Supporting Information Figure S1 for "Continuous B- to A-Transition in Protein-DNA Binding - How Well Is It Described by Current AMBER Force Fields?"

**Figure S1.** RMSD of simulated protein-DNA complexes for all three tested force fields separately for the complex, DNA and protein.

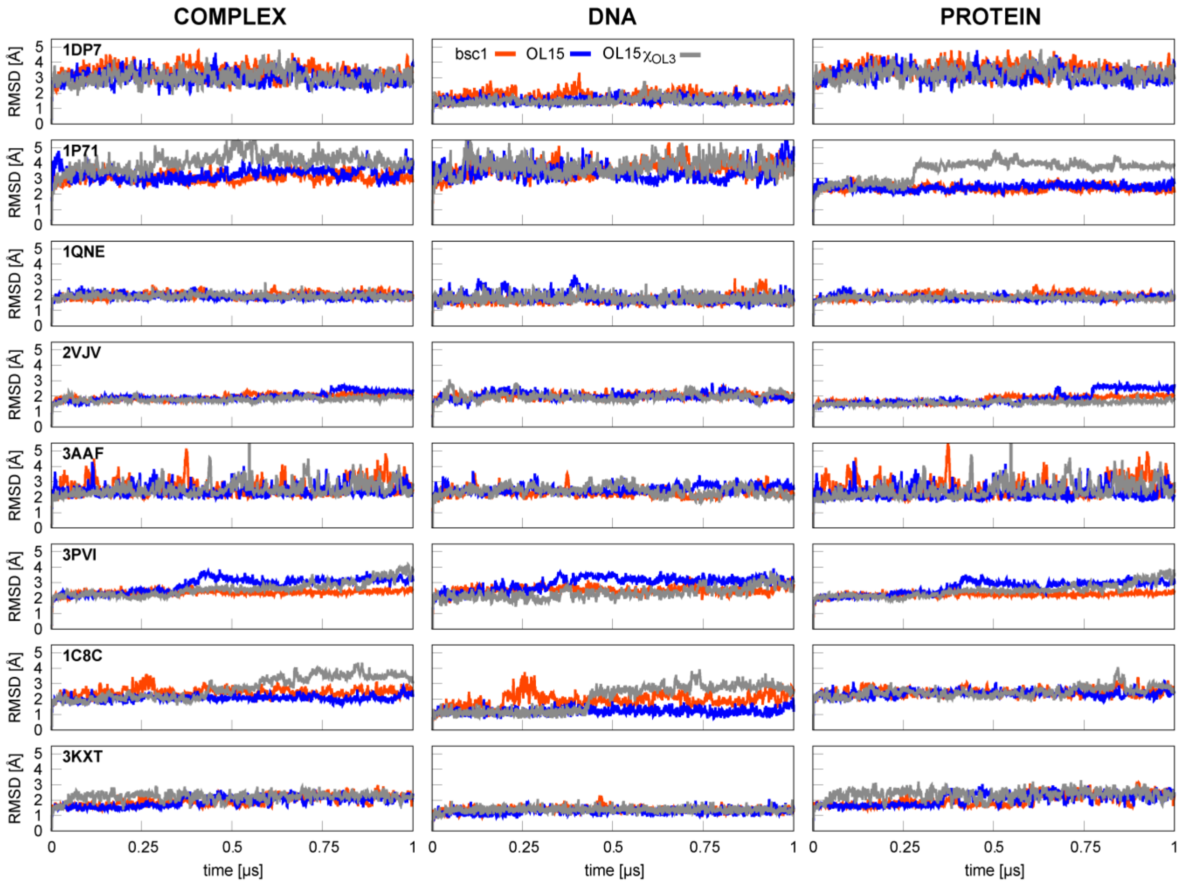


**Table S1.** The propensity to different P/χ states in complexes and in their naked DNAs averaged over all bsc1 simulations.

| Initial state in complex | % P/χ in complex | | | | % P/χ in naked DNA | | | |
|---|---|---|---|---|---|---|---|---|
| | A/A | A/B | B/A | B/B | A/A | A/B | B/A | B/B |
| A/A | 26.1 | 2.8 | 17.2 | 54.0 | 7.5 | 0.9 | 13.0 | 78.6 |
| A/B | 14.6 | 61.7 | 1.7 | 17.6 | 2.5 | 1.0 | 19.6 | 72.5 |
| B/A | 2.5 | 0.0 | 54.1 | 33.4 | 1.3 | 0.3 | 37.4 | 51.0 |
| B/B | 5.2 | 2.6 | 10.4 | 81.0 | 3.3 | 2.1 | 11.2 | 82.7 |

**Table S2.** The propensity to different P/χ states in complexes and in their naked DNAs averaged over all OL15$\chi_{OL3}$ simulations.

| Initial state in complex | % P/χ in complex | | | | % P/χ in naked DNA | | | |
|---|---|---|---|---|---|---|---|---|
| | A/A | A/B | B/A | B/B | A/A | A/B | B/A | B/B |
| A/A | 33.4 | 2.7 | 19.6 | 44.3 | 14.5 | 3.4 | 18.6 | 63.5 |
| A/B | 14.9 | 65.3 | 1.7 | 13.8 | 8.4 | 2.7 | 16.4 | 68.2 |
| B/A | 6.0 | 0.4 | 54.9 | 28.7 | 9.1 | 1.1 | 38.2 | 41.7 |
| B/B | 10.0 | 4.8 | 13.3 | 71.2 | 8.6 | 3.9 | 15.9 | 70.9 |
